# Generation and validation of a CRF_1_:Cre transgenic rat: The role of central amygdala CRF_1_ in nociception and anxiety-like behavior

**DOI:** 10.1101/2021.02.23.432551

**Authors:** Marcus M. Weera, Abigail E. Agoglia, Eliza Douglass, Zhiying Jiang, Shivakumar Rajamanickam, Rosetta S. Shackett, Melissa A. Herman, Nicholas J. Justice, Nicholas W. Gilpin

## Abstract

Corticotropin-releasing factor type-1 (CRF_1_) receptors are critical to stress responses because they allow neurons to respond to CRF released in response to stress. Our understanding of the precise role of CRF_1_-expressing neuronal populations in CRF-mediated behaviors has been largely limited to mouse experiments due to the lack of genetic tools available to selectively visualize and manipulate CRF_1_^+^ cells in rats. Here, we describe the generation and validation of a transgenic CRF_1_:Cre-^td^Tomato rat, which expresses a bicistronic *iCre-2A-^td^Tomato* transgene directed by 200kb of promoter and enhacer sequence surrounding the *Crhr1* cDNA present within a BAC clone, that has been transgenically inserted into the rat genome. We report that *Crhr1* and *Cre* mRNA expression are highly colocalized in CRF_1_:Cre-^td^Tomato rats within both the central amygdala (CeA), composed of mostly GABAergic neurons, and in the basolateral amygdala (BLA), composed of mostly glutamatergic neurons. In the CeA, membrane properties, inhibitory synaptic transmission, and responses to CRF bath application in ^td^Tomato^+^ neurons are similar to those previously reported in GFP^+^ cells in CRFR1-GFP mice. We show that stimulatory DREADD receptors can be selectively targeted to CeA CRF_1_^+^ cells via virally delivered Cre-dependent transgenes, that transfected Cre/^td^Tomato^+^ cells are activated by clozapine-n-oxide *in vitro* and *in vivo*, and that activation of these cells *in vivo* increases anxiety-like behavior and nocifensive responses. Given the accuracy of expression in the CRF_1_:Cre rat, modern genetic techniques used to investigate the anatomy, physiology, and behavioral function of CRF_1_^+^ neurons and circuits can now be performed in assays that require the use of rats as the model organism.

**Impact Statement:** A novel transgenic rat for studying the role of specific corticotropin-releasing factor type-1 receptor-expressing cell populations in physiology and behavior.

## Introduction

Corticotropin-releasing factor (CRF) is a 41 amino acid neuropeptide that acts on various cell populations in the brain and elicits stress-related behavioral (e.g., anxiety) and physiological (e.g. sympathomimetic) responses (Vale et al., 1981; Henckens et al., 2016). In the hypothalamic-pituitary-adrenal (HPA) axis, CRF released by neurons in the paraventricular nucleus of the hypothalamus (PVN) into the hypophyseal portal system stimulates adrenocorticotropic hormone (ACTH) release from the pituitary, which in turn stimulates release of glucocorticoids from the adrenal cortex. Glucocorticoids released into the circulatory system serve as effectors of the HPA axis by modulating various physiological (e.g., cardiovascular, respiratory, and immune) functions (Smith and Vale, 2006). Extrahypothalamic CRF neurons are located in many brain areas that modulate affective states and behaviors, such as the amygdala, cortex, hippocampus, midbrain, and locus coeruleus (Peng et al., 2017). Among these brain areas, the extended amygdala, particularly the central amygdala (CeA) and bed nucleus of stria terminalis (BNST), contain the highest densities of CRF neurons (Peng et al., 2017).

CRF type-1 (CRF_1_) receptors are G_s_- or G_q_-coupled metabotropic receptors (Milan-Lobo et al., 2009; Wanat et al., 2008) that are highly expressed in brain areas that contain high densities of CRF fibers and/or CRF neurons, including the CeA, BNST, and medial amygdala. CRF signaling via CRF_1_ modulates neurophysiological processes underlying stress reactivity, anxiety-related behaviors, learning and memory processes including fear acquisition and/or expression, pain signaling, and addiction-related behaviors (see reviews by Dedic et al., 2018, Henckens et al., 2016). Therefore, elucidation of the circuit-specific and cell-specific mechanisms by which CRF_1_ signaling mediates these processes represents an important avenue of research for understanding behaviors related to stress, anxiety, fear, pain, and addiction.

The central amygdala (CeA) is a key nucleus in modulating affective states and behaviors related to stress, anxiety, fear, pain, and addiction (Gilpin et al., 2015). Functionally, the CeA serves as a major output nucleus of the amygdala and can be anatomically divided into lateral (CeAl) and medial (CeAm) subdivisions (Duvarci and Pare, 2014). In mice and rats, CRF-expressing (CRF^+^) neurons are densely localized to the CeAl, whereas CRF_1_-expressing (CRF_1_^+^) neurons are mostly localized to the CeAm (Day et al., 1999; Jolkkonen and Pitkanen, 1998; Justice et al., 2008; Pomrenze et al., 2015). In addition to receiving CRF input from CRF^+^ neurons in the CeAl, the CeAm receives CRF inputs from distal brain areas such as the bed nucleus of stria terminalis (BNST) (Dabrowska et al., 2016) and dorsal raphe nucleus (Commons et al., 2003). CeA CRF_1_ signaling plays an important role in modulating affective states and behaviors. For example, pharmacological blockade of CeA CRF_1_ attenuates stress-induced increases in anxiety-like behavior (Henry et al., 2006), nociception (Itoga et al., 2016), and alcohol drinking (Roberto et al., 2010; Weera et al., 2020), as well as fear acquisition and expression (Sanford et al., 2017). Work using transgenic CRF and CRF_1_ reporter and Cre-driver mice have shed light on circuit- and cell-specific mechanisms by which CeA CRF^+^ and CRF_1_^+^ cells mediate affective behaviors (e.g., Fadok et al., 2017; Sanford et al., 2017). However, mice have a limited behavioral repertoire when compared to rats, making the study of more complex behaviors (e.g., drug self-administration and social interaction; Homberg et al., 2017) difficult.

Here, we describe the generation of a novel CRF_1_:Cre-^td^Tomato transgenic rat line. We show that *Crhr1* and *iCre* are expressed in the same neurons using hybridization histochemistry. In addition, we show that ^td^Tomato^+^ neurons are abundant in the CeAm, and that they are surrounded and in contact with CRF-containing puncta. We recorded membrane properties, inhibitory synaptic transmission, and spontaneous firing in CeA CRF_1_ cells and show that these cells are sensitive to CRF. In addition, we show that Cre-dependent DREADD receptors can be targeted for expression by CeA CRF_1_ cells such that DREADD stimulation of CRF_1_^+^ CeA neurons increases nociception and anxiety-like behaviors, recapitulating prior work using pharmacological strategies. These anatomical, electrophysiological and functional data support the utility of this CRF_1_:Cre-^td^Tomato transgenic rat line for the study of CRF_1_ neural circuit function, and will be an important new resource that will complement CRFR1:GFP and CRFR1:Cre mice (Justice et al., 2008; Jiang et al, 2018; Sanford et al., 2017), and CRF:Cre rats (Pomrenze et al., 2015) for the study of CRF signaling in physiology and behavior.

## Materials & Methods

| Key Resources Table |  |  |  |  |
| --- | --- | --- | --- | --- |
| Reagent type (species) or resource | Designation | Source or reference | Identifiers | Additional information |
| Gene (*Rattus norvegicus*) | BAC containing the *Crfr1* genomic locus | Riken Gene Engineering Division | RNB2-336H12 |  |
| Transfected construct | *iCre-2a-tdTomato* inserted BAC transgene | This paper |  |  |
| Genetic reagent (*Rattus norvegicus*) | CRF1:Cre-<sup>td</sup>Tomato rat | This paper |  |  |
| Sequence-based reagent |  |  |  |  |
| Recombinant DNA reagent | AAV8-hSyn-DIO-HA-hM3D(Gq)-IRES-mCitrine | Addgene | Cat# 50454-AAV8 |  |
| Recombinant DNA reagent | AAV5-hSyn-DIO-EGFP | Addgene | Cat# 50457-AAV5 |  |
| Antibody | Anti-c-Fos (Rabbit polyclonal) | Abcam | Cat# ab190289, RRID:AB_2737414 | 1:1000 |
| Antibody | Anti-HA-Tag (Rabbit monoclonal) | Cell Signaling | Cat# 3724, RRID:AB_1549585 | 1:250 |
| Antibody | Anti-RFP (Rabbit monoclonal) | Abcam | Cat# ab34771, RRID:AB_777699 | 1:500 |
| Antibody | Anti-CRF (Rabbit monoclonal) | The Salk Institute | Rc-68 | 1:2000 |
| Commercial assay or kit | RNAscope Multiplex Fluorescent Kit v2 | ACD Bio | *iCre* and *Crfr1* probes |  |
| Commercial assay or kit | TSA Detection Kit | Akoya Biosciences | Cat# NEL701A001KT |  |
| Chemical compound, drug | Clozapine-n-oxide | NIH Drug Supply Program |  |  |
| Chemical compound, drug | Corticotropin-releasing factor | Tocris | Cat# 1607 |  |
| Software, algorithm | Mini Analysis | Synaptosoft Inc. | RRID:SCR_002184 |  |
| Software, algorithm | Clampfit 10.6 | Molecular Devices |  |  |
| Software, algorithm | Prism 7.0 | GraphPad | RRID:SCR_002798 |  |
| Software, algorithm | SPSS 25 | IBM SPSS | RRID:SCR_019096 |  |
| Other | Prolong Gold Antifade Reagent with DAPI | Invitrogen | Cat# P36935 |  |

### Development of CRF_1_:Cre-^td^Tomato rats

#### Design of CRF_1_:Cre bacterial artificial chromosome (BAC) transgene

Please refer to **Figure 1** for a schematic of the BAC generation. The design of the CRF_1_:Cre rat BAC transgene is similar to the design used to generate CRFR1:Cre BAC transgenic mice (Jiang et al., 2018), with the exception that a BAC clone from rat was used (clone RNB2-336H12 from Riken Rat genomic BAC library). The BAC clone used lacks any other identified protein encoding genetic sequences reducing the possibility that other genes will be expressed from BAC when introduced into the rat genome.. We obtained the RNB2-336H12 clone from the Riken Rat BAC clone collection (Saar et al., 2008).This BAC was inserted using recombineering techniques such that sequences encoding *Crhr1* were replaced, beginning at the translation ATG start site, with seuences endocing a bicistronic *iCre-2A-^td^Tomato* encoding transgene (**Fig. 1C**). *iCre* is a codon-optimized Cre that produces consistent Cre activity (Koresawa et al., 2000). ^td^Tomato encodes a red fluorescent protein that allows cells expressing the transgene to be visualized (Shaner et al., 2004). The poly A and WPRE sequences stabilize the mRNA to achieve more robust expression (Glover et al., 2002). To insert the iCre-p2A-tdTomato cassette into the BAC, we transformed bacterial cells that contain the BAC with a helper plasmid (Portmage-4) which contains a heatshock inducible element that drives expression of lambda red recombinase and confers chloramphenicol resistance (Liu et al., 2003). These cells were then transformed with the iCre-p2A-^td^Tomato homology arm targeting cassette, after a 15-min heatshock. Cells were selected on kanamycin (for neoR) and colonies were screened by colony PCR for accurate insertion of the cassette. Confirmed inserted BAC DNA was then purified transformed into EL250 cells that contained an arabinose inducible flipase construct that were grown in L-arabinose for 1 hr. The transformation was selected on ampicillin (the resistance of the BAC) and screened for loss of kanamycin resistance. Colonies were screened with original (5’) and new (flipped out neo) PCR products. Bacteria containing the final full-length inserted BAC construct were sent to the University of North Carolina (UNC) Transgenic Core, where the BAC DNA was expanded and purified, checked for integrity by DNA laddering with EcoR1 and XbaI and pulsed-gel electrophoresis as well as PCR, linearized, then injected into single celled rat oocytes.

**Figure 1.**
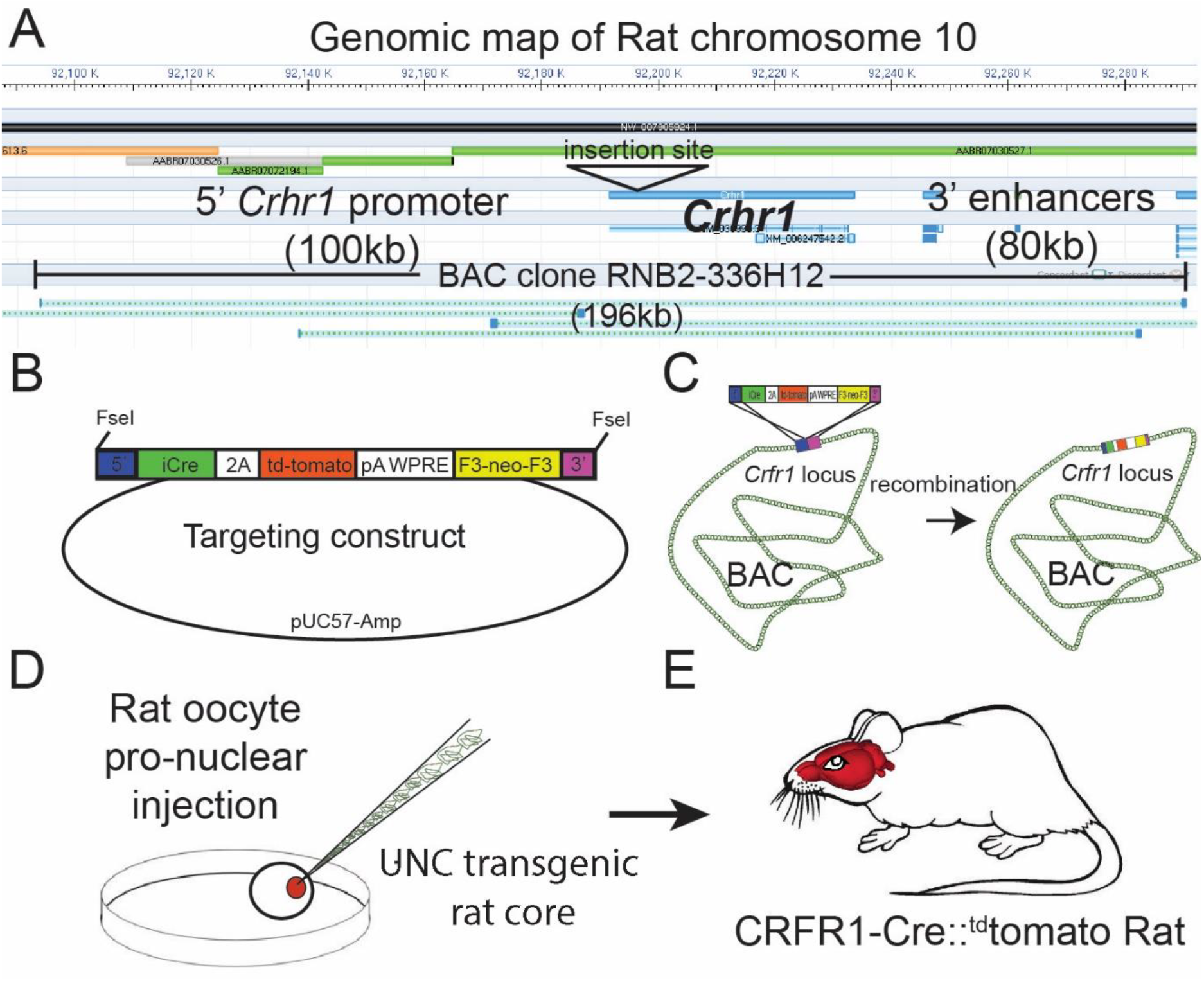
Design of the Crhr1-Cre2aTom BAC transgene. **(A)** *Crhr1* is located on chromosome 10 in the rat. A BAC clone containing 196kb of DNA surrounding the *Crhr1* coding region includes 100kb of upstream and 80kb of downstream DNA, where the majority of promoter and enhancer sequences that control Crhr1 expression are located, was obtained (Riken, RNB2-336H12). There are no other sequences within this 196kb DNA clone have been annotated as coding sequences for genes other than Crhr1. **(B)** A transgene containing 5’ (blue) and 3’ (magenta) targeting sequences, a bicistronic iCre 2A fused tdTomato (red) sequence, 3’ polyA/WPRE stabilizing sequence, and a F3 flanked neomycin resistance sequence (yellow) was constructed then transformed into E.Coli containing the RNB2-336H12 BAC construct. **(C)** Using recombineering techniques we isolated BAC clones in which targeted insertion of the transgene at the translation start site of *Crfr1* (ATG) was confirmed by PCR/sequencing. A single Bacterial clone containing the transgene inserted BAC was sent to the UNC transgenic facility where BAC DNA was purified and injected into single cell, fertilized rat oocytes. Two independent rat lines were recovered in which the entire BAC sequence (confirmed by PCR) was inserted into genomic DNA, of which one line displayed transgenic expression in a pattern representative of known Crhr1 expression patterns.

#### Generation of transgenic rats

Transgenic rats were generated by the UNC Transgenic Core by injectingsingle-cell rat oocytes with modifiedBAC (**Fig 1D, E**). DNA from F1 offspring were tested for the presence of the introduced BAC transgene using PCR. Two animals containing BAC insertions were further tested for the presence of BAC vector specific sequences from both BAC termini and found to be positive, indicating the BAC inserted contained the entire BAC sequence. Transgenic rats were first outcrossed to wildtype Wistar animals, then intercrossed to maximize the genetic similarity of transgenic offspring. Transgenic animals were generated on a Wistar backgroud because Wistar rats are commonly used in models of alcohol and substance use disorders, and other models of behavioral and physiological disorders.

#### Subjects for experiments

Adult male and female Crhr1:Cre-^td^Tomato rats were used in all experiments. Rats bred from the original founder F1 line were group-housed in humidity- and temperature-controlled (22°C) vivaria at UNC, LSUHSC, or UTHSC on a reverse 12 h light/dark cycle (lights off at 7:00 or 8:00 AM) and had *ad libitum* access to food and water. All behavioral procedures occurred in the dark phase of the light-dark cycle. Sample sizes were estimated using published work from our labs. *In situ* hybridization and immunohistochemistry experiments were performed in a single pass. Behavioral experiments were performed in two independent, fully counterbalanced replicates. Slice electrophysiology experiments were conducted in a single pass, such that one experiment was performed in each slice and each experimental group contained neurons from a minimum of 3 rats. All procedures were approved by the Institutional Animal Care and Use Committee of the respective institutions at which procedures occurred and were in accordance with National Institutes of Health guidelines.

### Stereotaxic surgeries

Rats were anesthetized with isoflurane (2 min at 4% for induction, 1-3% for maintenance) and mounted into a stereotaxic frame (Kopf Instruments) for all stereotaxic surgeries. The following coordinates (from bregma) were used for bilateral intra-CeA microinjections: −2.5 mm posterior, ± 4.0 mm lateral, and −8.4 mm ventral for male rats and −2.2 mm posterior, ± 4.0 mm lateral, and − 8.2 mm ventral for female rats. Viral vectors for Cre-dependent expression of Gq-DREADDs or control (see below) were injected into each side of the CeA at a volume of 0.5 μL over 5 min and injectors were left in place for an additional 2 min. Viral titers were between 1.0-1.5 x 10^13^ GC/mL. At the end of surgeries, rats were monitored to ensure recovery from anesthesia and were given a minimum of 4 weeks to recover before the start of procedures. Rats were treated with the analgesic flunixin (2.5 mg/kg, s.c.) and, in some rats, the antibiotic cefazolin (20 mg/kg, i.m.) before the start of surgeries and once the following day.

### Immunohistochemistry

Rats were deeply anesthetized with isoflurane and were transcardially perfused with ice-cold phosphate buffered saline (PBS) followed by 4% paraformaldehyde (PFA). Brains were extracted and post-fixed in 4% PFA for 24 h (at 4°C), cryoprotected in 20% sucrose for 48-72 h (at 4°C), and frozen in 2-methylbutane on dry ice. Coronal sections were collected using a cryostat and stored in 0.1% sodium azide in PBS at 4°C until further processing.

#### c-Fos immunofluorescent labeling

Sections (40 μm) containing the CeA were washed 3 x 10 min in PBS and incubated in blocking buffer (2.5% normal goat serum with 0.3% Triton X-100) for 2 hr at RT. Subsequently, sections were incubated in rabbit anti-c-Fos polyclonal antibody (1:1000 in blocking buffer; catalog no. 190289, Abcam, Cambridge, United Kingdom) for 48 hr at 4°C. Sections were then washed 3 x 10 min in PBS and incubated in goat anti-rabbit Alexa Fluor 647 (1:500 in blocking buffer; catalog no. A32733, Invitrogen, Carlsbad, CA) for 2 hr at RT. After 3 x 10 min washes in PBS, sections were mounted on microscope slides and coverslipped with Prolong Gold Antifade Reagent with DAPI (Invitrogen, catalog no. P36935). Sections were imaged using a Keyence (Osaka, Japan) BZ-X800 fluorescent microscope at 20x magnification and Fos^+^ cells were quantified manually by a blinded experimenter. Four sections representative of the CeA anterior-posterior axis (~bregma −1.8 to −2.8) from each animal were used for analysis.

#### HA-tag immunofluorescent labeling

Sections (40 μm) containing the CeA were washed 3 x 10 min in PBS and incubated in 3% hydrogen peroxide for 5 min. Sections were then washed 3 x 10 min in PBS and incubated in a blocking buffer containing 1% (w/v) bovine serum albumin and 0.3% Triton X-100 in PBS for 1 h at room temperature (RT). Then, sections were incubated in a rabbit anti-HA monoclonal antibody (1:250 in blocking buffer; catalog no. 3724, Cell Signaling, Danvers, MA) for 48 h at 4°C. Sections were then washed for 10 min in TNT buffer (0.1 M Tris base in saline with 0.3% Triton X-100), incubated for 30 min in 0.5% (w/v) Tyramide Signal Amplification (TSA) blocking reagent in 0.1 M Tris base, and incubated for 30 min in ImmPRESS horseradish peroxidase horse anti-rabbit antibody (catalog no. MP-7401, Vector Laboratories, Burlingame, CA) at RT. Following 4 x 5 min washes in TNT buffer, sections were incubated in fluorescein TSA reagent (1:50 in TSA amplification diluent) for 10 min at RT. TSA blocking reagent, fluorescein TSA reagent, and TSA amplification diluent are part of the TSA detection kit (catalog no. NEL701A001KT, Akoya Biosciences, Marlborough, MA). Sections were washed 3 x 10 min in TNT buffer, mounted on microscope slides, and coverslipped with Prolong Gold Antifade Reagent with DAPI (Invitrogen, catalog no. P36935). Sections were imaged using a Keyence BZ-X800 fluorescent microscope at 2x and 20x magnification.

#### ^td^Tomato DAB immunostaining

Because ^td^Tomato signal degrades over time, to create a permanent set of slides for anatomical analysis we performed immunohistochemistry to label ^td^Tomato protein permanently. Briefly, fixed, free-floating sections (30 μm) were incubated overnight in monoclonal rabbit anti-RFP, biotin tagged antibody (1:500; Abcam, catalog no. ab34771). Sections were then washed 3x with PBS and incubated in streptavidin-conjugated peroxidase (DAB-elite kit) for 1 hr. After incubation with streptavidin, sections were washed 2x in PBS, then 2x in 0.1M NaOAc (pH 6.0), then stained in a solution of 0.1M NaOAC (pH 6.0) containing nickel ammonium sulfate (3%) and 5 μl of 3% H_2_0_2_. Sections were stained for up to 10 min, then washed 2x in NaOAc (pH 6.0), then in PBS, before being mounted on gelatin coated slides, dehydrated, and coverslipped in DPX. Bright field images were acquired using a Cytation 5 imager (BioTek Instruments, Winooski, VT).

#### ^td^domato and CRF immunofluorescent labeling

Tissue processing procedures were similar to the immunofluorescent procedures described above. Sections through the amygdala were incubated in antibodies against CRF (rabbit anti-CRF, # rc-68, 1:2000, The Salk Institute) and goat-anti-RFP (1:1000; Rockland, catalog no. 200-101-379). Primary antibodies were detected by secondary anti-rabbit antibody conjugated with Alexa Fluor 488 and anti-goat antibody conjugated with Alexa Fluor 555 (Invitrogen), resulting in ^td^Tomato protein being visible as red fluorescence and CRF peptide visible as green fluorescence. High magnification images of the CeA were taken using a Leica (Wetzlar, Germany) Sp5 confocal microscope.

### *In situ* hybridization

All solutions were prepared with DEPC treated water and all tools and surfaces were wiped with RNAzap followed by DEPC treated water. Adult Crhr1:Cre −^td^Tomato rats (3 males and 3 females) were deeply anesthetized with Avertin (2,2,2,-tribromoethanol, 1.25% solution, 0.2 ml/10 g BW, IP), then transcardially perfused with PBS followed by 4% PFA in PBS. Brains were removed, fixed in 4% PFA at 4°C overnight, then equilibrated in 30% sucrose, sectioned into six series of sections (30 μM, coronal sections) on a frozen sliding microtome (SM 2000R, Leica), and stored in PBS at 4°C. Brain slices were mounted onto glass slides, dried, and went through *in situ* hybridization (ISH) using a RNAscope Multiplex Fluorescent kit v2 (ACDbio, Newark, CA) following the manufacturer’s protocol.

### Slice electrophysiology

Following rapid decapitation, brains were extracted and sectioned as previously described (Herman & Roberto, 2016). Briefly, brains were placed in ice-cold high sucrose solution containing (in mM): sucrose 206.0; KCl 2.5; CaCl2 0.5; MgCl2 7.0; NaH2PO4 1.2; NaHCO3 26; glucose 5.0; HEPES 5. Coronal sections (300μm) were prepared on a vibrating microtome (Leica VT1000S, Leica Microsystem) and placed in an incubation chamber with oxygenated (95% O2/5% CO2) artificial cerebrospinal fluid (aCSF) containing (in mM): NaCl 120; KCl 2.5; EGTA 5; CaCl2 2.0 MgCl2 1.0; NaH2PO4 1.2; NaHCO3 26; Glucose 1.75; HEPES 5. Slices were incubated for 30 min at 37 °C, followed by a 30 min acclimation at room temperature. Patch pipettes (3-6 MΏ; King Precision Glass Inc., Claremont, CA) were filled with an internal solution containing (in mM): potassium chloride (KCl) 145; EGTA 5; MgCl2 5; HEPES 10; Na-ATP 2; Na-GTP 0.2 (for whole cell voltage-clamp recordings) or containing potassium gluconate 145; EGTA 5; MgCl2 5; HEPES 10; Na-ATP 2; Na-GTP 0.2 (for whole cell current clamp experimental recordings). Data acquisition was performed with a Multiclamp 700B amplifier (Molecular Devices, San Jose, CA), low-pass filtered at 2-5 kHz, coupled to a digitizer (Digidata 1550B; Molecular Devices) and stored on a PC using pClamp 10 software (Molecular Devices). Whole-cell voltage-clamp recordings were performed at Vhold = −60 mV. Cell-attached recordings were performed with no holding parameters (0 mV). All recordings were performed at room temperature. Series resistance was continuously monitored and cells with series resistance >15 MΩ were excluded from analysis. Properties of sIPSCs were analyzed and visually confirmed using a semi-automated threshold detection program (Minianalysis). The frequency of firing discharge was evaluated and visually confirmed using threshold-based event detection analysis in Clampfit 10.2 (Molecular Devices). Experimental drugs were applied by bath application or y tube for focal application. Analysis was performed on recordings containing >60 events or that encompassed a period of 2-5 minutes.

### Drugs

Clozapine-n-oxide (CNO, NIH Drug Supply Program) was dissolved in 5% DMSO (v/v in saline). CRF was purchased from Tocris Bioscience (Bristol, United Kingdom), dissolved in stock solutions in ultra-pure water, and diluted to final experimental concentration in aCSF.

### Experiment 1: Crhr1 and iCre expression in CeA

The purpose of this experiment was to determine the pattern of ^td^Tomato protein expression and *Crhr1* and *iCre* mRNA expression within the CeA. Coronal brain sections containing the CeA were processed for immunohistochemical DAB labeling of ^td^Tomato protein or RNAscope ISH for labeling *Crhr1* and *iCre* mRNA, as described above. A separate set of brain sections containing CeA were processed for immunofluorescent labeling of ^td^Tomato and CRF to map the expression pattern of these proteins in the CeA of Crhr1:Cre-^td^Tomato rats. Images were captured using a confocal microscope (model TCS SP5, Leica) and processed with Fiji ImageJ. Coronal sections containing the CeA were identified by neuroanatomical landmarks with reference to a rat brain atlas and captured at 20x magnification at one single focal plane (1 μm). For analysis of *Crhr1* and *iCre* RNAscope images, punctate signals from each channel were quantified separately following the manufacturer’s guideline (ACDbio SOP45-006). Quantification was performed by an experimenter blinded to experimental groups. Based on pilot studies, cells that had more than 3 puncta were considered positive. At least 3 sections representative of the anterior-posterior axis of the CeA were analyzed in each animal.

### Experiment 2: Membrane properties, inhibitory synaptic transmission, and CRF sensitivity of CeA CRF_1_:Cre-^td^Tomato cells

The purpose of this experiment was to characterize intrinsic properties, inhibitory synaptic transmission, and CRF sensitivity in CRF_1_^+^ neurons in the CeA. These neurons were identified using fluorescent optics and brief (<2 s) episcopic illumination. All labeled neurons were photographed, recorded, and saved. Intrinsic membrane properties were determined in voltage clamp configuration (V_hold_ = −60mV) using pClamp 10 Clampex software. Current clamp recordings were performed to determine current-voltage (I-V) changes and the firing type of each neuron. Voltage clamp recordings of pharmacologically-isolated GABA_A_ receptor-mediated spontaneous inhibitory postsynaptic currents (sIPSCs) were performed with bath application of the glutamate receptor blockers 6,7-dinitroquinoxaline-2,3-dione (DNQX, 20μM) and DL-2-amino-5-phosphonovalerate (AP-5, 50 μM) and the GABAB receptor antagonist GCP55845A (1μM). Cell-attached recordings were made in close juxtacellular (i.e., membrane intact) cell-attached configuration and only cells with regular spontaneous firing were included in analysis. After a stable baseline period, CRF (200 nM) was applied for a period of 7-10 min and changes in firing were measured and compared to baseline. Experiments were performed in individual slices to ensure that drug application was never repeated in the same slice.

### Experiment 3: Functional validation of Cre-dependent expression of Gq-DREADD receptors in CeA CRF_1_:Cre -^td^Tomato cells

The purpose of this experiment was to test if CeA CRF_1_:Cre-^td^Tomato cells can be activated using chemogenetics via Cre-mediated targeting of Gq-DREADD receptors [hM3D(Gq)] to these cells.

#### c-Fos validation

To test if Cre-dependent expression of Gq-DREADD receptors can stimulate CeA CRF_1_:Cre-^td^Tomato cells, CRF_1_:Cre-^td^Tomato rats were given bilateral microinjections of AAV8-hSyn-DIO-HA-hM3D(Gq)-IRES-mCitrine (50454-AAV8, Addgene, Watertown, MA) or a control virus (AAV5-hSyn-DIO-EGFP; Addgene, 50457-AAV5) targeting the CeA. Four weeks later, rats were given a systemic CNO injection (4 mg/kg, i.p.) and sacrificed 90 min later. Brain sections (4 sections/rat x 4 rats/group) were processed for c-Fos immunohistochemistry and the percentage of c-Fos^+ td^Tomato^+^ cells in the CeA was calculated. Cell counts were performed by an experimenter blinded to experimental groups.

#### Electrophysiological validation

To functionally validate Cre-dependent expression of Gq-DREADD receptors in CeA CRF_1_:Cre -^td^Tomato cells, CRF_1_:Cre-^td^Tomato rats were given bilateral microinjections as described above. After a minimum of four weeks, brain slices of CeA were prepared as described above. CeA neuronal expression of mCitrine and tdTomato were confirmed by fluorescent optics and neurons were targeted for electrophysiological recording. After a stable baseline period, CNO (10 μM) was applied and changes in membrane potential and action potential firing were measured and compared to baseline.

### Experiment 4: Effects of chemogenetic stimulation of CeA CRF_1_:Cre -^td^Tomato cells on nociception and anxiety-like behavior

The purpose of these experiments was to test the effects of chemogenetic stimulation of CeA CRF_1_ cells on nociception and anxiety-like behavior. Rats were given intra-CeA microinjections of AAV8-hSyn-DIO-HA-hM3D(Gq)-IRES-mCitrine (Addgene, 50454-AAV8; Active Virus) or AAV5-hSyn-DIO-EGFP (Addgene, 50457-AAV5; Control Virus) and were given 4 weeks for recovery and viral expression (**Fig. 7A**). Please refer to **Figure 7B** for a timeline schematic of this experiment. All rats were habituated to handling before the start of behavioral procedures. On behavioral procedure days, rats were given at least 30 min to acclimate to the procedure room.

#### Nociception

Mechanical and thermal nociception were measured using the Von Frey (Pahng et al., 2017) and Hargreaves (Avegno et al., 2018) assays, respectively, as previously described. Briefly, the Von Frey apparatus consists of clear chambers placed on top of a mesh floor. To measure sensitivity to mechanical nociception, each hind paw was perpendicularly stimulated with a Von Frey filament (Electronic Von Frey 38450, Ugo Basile, Gemonio, Italy) calibrated to measure the amount of force applied using the up-down method and the force (g) threshold required to elicit a paw withdrawal response was recorded. Force thresholds were measured twice for each hind paw in alternating fashion, with at least 1 min between measurements, and an average threshold was calculated for each animal. The Hargreaves apparatus consists of clear chambers placed on top of a glass pane suspended above a tabletop. To measure sensitivity to thermal nociception, each hind paw was stimulated by a halogen light heat source (Model 309 Hargreaves Apparatus, IITC Life Sciences, Woodland Hills, CA) and latency (s) for hind paw withdrawal was measured. Withdrawal latencies were measured twice for each hind paw in alternating fashion, with at least 1 min between measurements, and an average withdrawal latency was calculated for each animal.

Baseline paw withdrawal thresholds in the Von Frey assay and withdrawal latencies in the Hargreaves assay were measured over 3 sessions (1 baseline session/day; baseline sessions for each assay occurred on alternating days; **Fig. 7B**) that were each preceded 30 min earlier by a vehicle (5% DMSO in saline, i.p.) pretreatment. After the final baseline session, rats were counterbalanced into CNO (4 mg/kg) or Vehicle (5% DMSO) treatment groups based on paw withdrawal latencies during the 3^rd^ (final) Hargreaves baseline session. During Von Frey and Hargreaves test sessions, rats were given CNO (4 mg/kg) or vehicle injections (i.p.) 30 min before the start of testing. All Von Frey and Hargreaves procedures occurred under regular white light illumination.

#### Anxiety-like behaviors

One day after Hargreaves testing, rats were tested for anxiety-like behaviors in the elevated plus maze (EPM), open field (OF), and light-dark box (LD) on consecutive days (**Fig. 7B**). All procedures occurred under indirect, dim illumination (~10 lux). Rats were given CNO (4 mg/kg) or vehicle injections (i.p.) 30 min before the start of each test. The EPM and OF tests were performed as previously described (Albrechet-Souza et al., 2020; Fucich et al., 2020). Briefly, the EPM consists of two open and two closed arms elevated 50 cm above the floor. Rats were individually placed in the center of the maze facing an open arm and were given 5 min to explore the maze. Time spent in the open and closed arms of the maze was measured. The OF consists of a square arena with a checkerboard patterned floor (4 x 4 squares). Rats were individually placed in one corner of the arena and were given 5 min to explore the arena. Time spent in the periphery and the center of the arena (defined as the 3 x 3 squares in the center of the arena) was measured. The LD box consists of a two compartments; one with black walls and a black floor, and the other with white walls and a white floor. The black compartment was protected from light (dark box) and the white compartment was illuminated (light box; ~1000 lux). Rats were able to freely explore both dark and light boxes through an opened door. Rats were individually placed in the dark box and were given 5 min to explore the apparatus. Time spent in dark and light boxes, as well as the latency to enter the light box were measured. EPM, OF, and LD tests were recorded via a camera mounted above the apparatus and videos were scored by an experimenter blinded to treatment groups. At the end of the experiment, rats were sacrificed and brain sections were analyzed for virus placement.

### Experiment 5:*Crhr1* and *iCre* expression in BLA

The purpose of this experiment was to examine *Crhr1* and *iCre* mRNA expression in a brain area outside the CeA. The basolateral amygdala (BLA) was selected for analysis because CRF_1_ receptors are highly expressed in, and functionally regulate the activity of pyramidal glutamatergic neurons in this area (as opposed to GABAergic neurons in CeA) (Refojo et al., 2011; Rostkowski et al., 2013). Brain sections containing BLA were processed for *Crhr1* and *iCre* RNAscope and analyzed as described in Experiment 1.

### Statistical analyses

Electrophysiology data on frequency and amplitude of spontaneous inhibitory postsynaptic currents (sIPSCs) were analyzed and manually confirmed using a semi-automated threshold-based detection software (Mini Analysis, Synaptosoft Inc., Decatur, GA). Cell-attached firing discharge data were analyzed and manually confirmed using a semi-automated threshold-based detection software (Clampfit 10.6, Molecular Devices). Electrophysiological characteristics were determined from baseline and experimental drug application containing a minimum of 65 events each. Event data were represented as mean ± SEM or mean % change from baseline ± SEM and analyzed for independent significance using a one-sample t-test, compared by paired or unpaired t-test where appropriate. Data analysis and visualization were completed using Prism 7.0 (GraphPad, San Diego, CA). Behavioral data were analyzed using multifactorial repeated measures ANOVAs (for Von Frey and Hargreaves tests) or t-tests (for anxiety tests). For RM ANOVAs, between-subjects factors include sex and treatment, and the within-subjects repeated measure was test session (i.e., baseline vs. test). Data from the active and control virus groups were analyzed separately (i.e., the control virus group was treated as a replication of the experiment; Weera et al., 2021). Data from experiments that only have two groups (e.g., c-Fos immunohistochemistry) were analyzed using t-tests. Data were analyzed using the Statistical Package for Social Sciences (Version 25, IBM Corporation, Armonk, NY). Statistical significance was set at *p* < 0.05.

## Results

### Experiment 1: iCre (^td^Tomato) is expressed in CRF_1_-expressing cells in the CeA

The purpose of this experiment was to determine the pattern of ^td^Tomato protein expression and *Crhr1* and *iCre* mRNA expression within the CeA. Immunohistochemical labeling of ^td^Tomato in the amygdala showed that ^td^Tomato^+^ cells were located in the lateral, basolateral, central, and medial amygdala (**Fig. 2A**). Within the CeA, ^td^Tomato^+^ cells were found to be concentrated in the CeAm, whereas the CeAl was largely devoid of ^td^Tomato^+^ cells (**Fig. 2B**).

**Figure 2.**
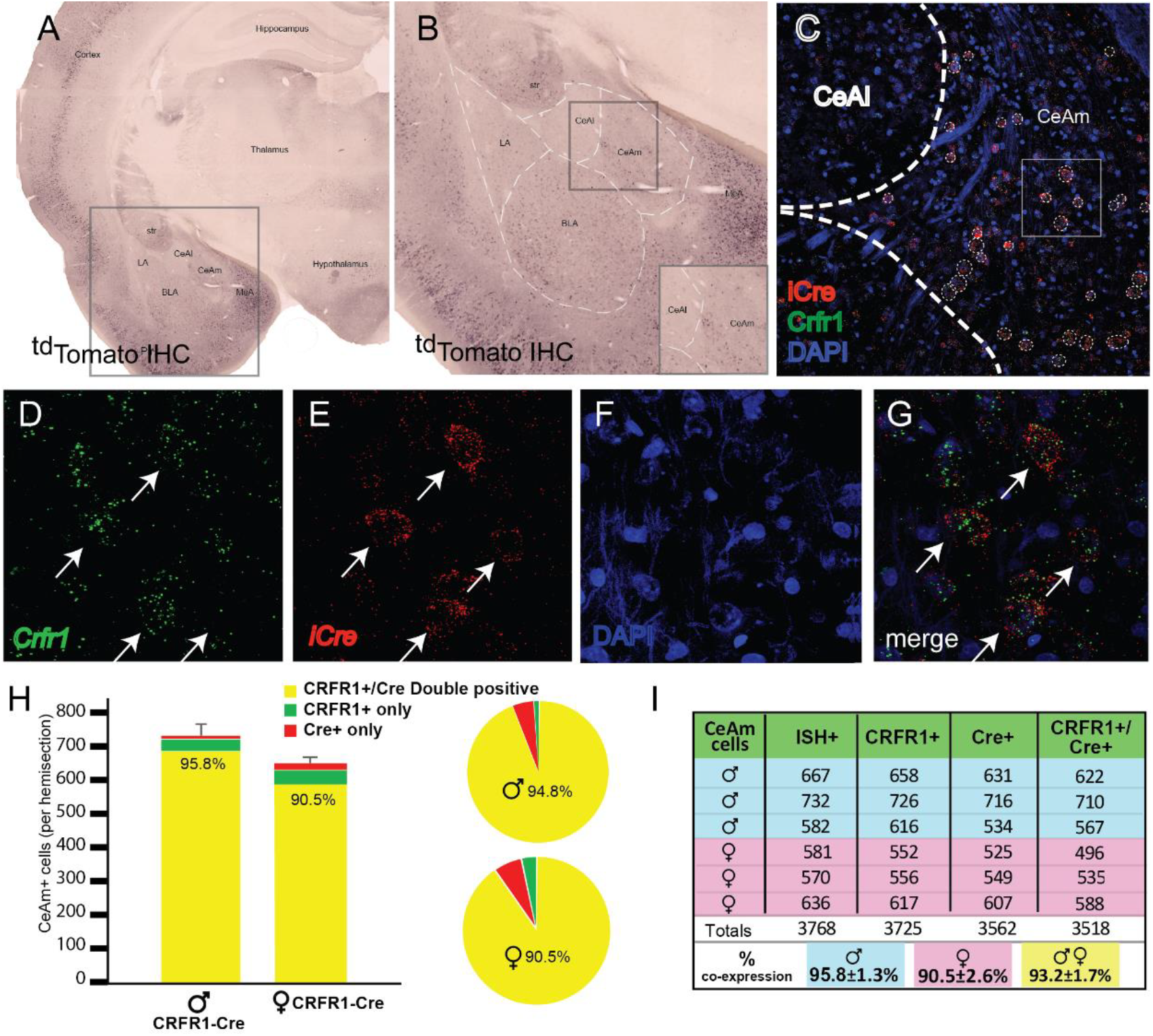
Validation of transgenic expression of iCre/^td^Tomato in CRF_1_^+^ expressing neurons located in the medial central nucleus of the amygdala (CeAm). **(A)** A low magnification image of a section containing the amygdala from a CRF_1_:Cre rat, immunohistochemically labeled for ^td^Tomato. Expression of the CRF_1_:Cre transgene is broadly very similar to previous reports of CRF_1_ expression in both rat and mouse. **(B)** Within the boxed region of panel A, higher magnification reveals CRF_1_^+^ cells in the lateral amygdala (LA), basolateral amygdala (BLA), medial portion central amygdala (CeAm), and medial amygdala (MeA). The lack of significant labeling in the CeAl is consistent with reports using both *in situ* hybridization and transgenic reporters to detect CRF_1_ expression. **(C)** Micrograph of the CeA from the region boxed in panel B allows visualization of mRNA encoding *iCre* (red) and *Crhr1* (green) along with nuclei stained with DAPI (blue). **(D-G)** Higher magnification images of the boxed region in (C) allows visualization of mRNA for *Crhr1* (D, green), and *iCre* (E, red), with nuclei visualized by DAPI staining (F). **(G)** Merged images reveals that many CeAm neurons that are positive for *Crhr1* mRNA are also positive for *iCre* mRNA (arrows point to double positive neurons). Quantification of coincidence of *in situ* hybridization for both *Crhr1* and *iCre* mRNAs demonstrates that >90% of *Crhr1* positive cells are also positive for *iCre* in the CeAm (n=3). **(H)** Graphical representation of quantification of coincident labeling, or **(I)** a table of the precise counts from each of 3 male and 3 female CRF_1_:Cre transgenic animals. We observed greater than 90% of neurons positive for both *Crhr1* and *iCre* in the CeAm.

We used RNAscope ISH to test the hypothesis that *Crhr1* and *iCre* mRNA are highly colocalized within the CeA. ^td^Tomato mRNA was not probed because iCre and ^td^Tomato are expressed as a single polypeptide (iCre-2A-^td^Tomato) that is cleaved at the 2A site. RNAscope ISH probing of *Crhr1* and *iCre* mRNA showed strong expression of these molecules within the CeAm (**Fig. 2C-G**). Since previous work (e.g., Justice et al., 2008) and our data show that CRF_1_^+^ cells are largely localized to the CeAm, analysis of *Crhr1* and *iCre* mRNA expression was focused on this subregion. Quantification of *Crhr1*- and *iCre*-expressing cells within the CeAm showed that more than 90% of *Crhr1*-expressing cells co-express *iCre* (**Fig. 2H, I**). There were no significant sex differences in the number of *Crhr1^+^*, *Cre^+^*, and *Crhr1^+^/Cre^+^* cells, but there was a trend for more *Crhr1*^+^ cells in the CeAm of male rats (*p* = 0.07).

Previous work showed that CeA CRF^+^ cells are concentrated in the CeAl, whereas CRF_1_^+^ cells are mostly located in the CeAm (Day et al., 1999; Jolkkonen and Pitkanen, 1998; Justice et al., 2008; Pomrenze et al., 2015). Immunofluorescent labeling of CRF and ^td^Tomato protein in the CeA of CRF_1_:Cre-^td^Tomato rats shows that the topography of CRF^+^ and ^td^Tomato^+^ (CRF_1_:Cre) cells in these rats is consistent with previous studies (**Fig. 3**).

**Figure 3.**
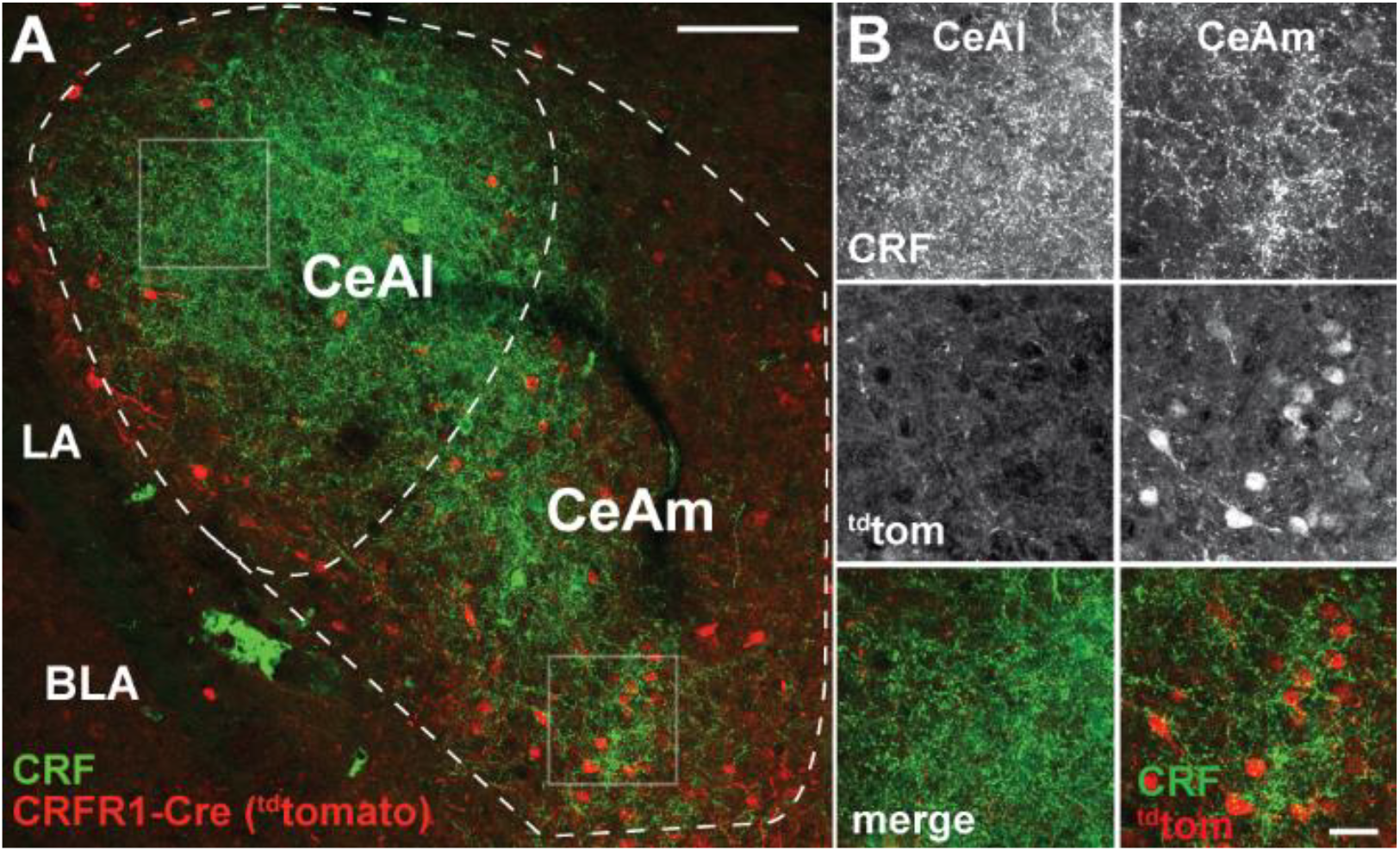
CRF_1_ driven expression of Cre/^td^Tomato in the CeA. **(A)** Visualization of CRF using immunofluorescent labeling (green) in a rat carrying the CRF_1_-Cre2aTom transgene (red) reveals minimal cellular expression of CRF_1_ in the lateral central nucleus of the amygdala (CeAl) where CRF is highly abundant. This discrepancy in CRF localization compared to CRF_1_ expression is consistent with previous reports of CRF_1_ expression in both rat and mouse. In contrast to the CeAl, the medial central nucleus of the amygdala (CeAm) contains many CRF_1_^+^ neurons (reported by the CRF_1_-Cre2Atom transgene), in contact with puncta positive for CRF peptide. **(B)** High resolution images from the boxed regions of CeAl and CeAm in panel A. CRF staining is dense in both the CeAl and CeAm (top panels), however cellular expression of the CRF_1_-Cre2aTom transgene is low in the CeAl, while many neurons in the CeAm are positive for CRF_1_ expression (middle panels). Merged images (lower panels) display the coincident staining of CRF_1_^+^neurons with CRF puncta in the CeAm, suggesting that stress driven CRF release directly signals to CRF_1_^+^ neurons in the CeAm to modulate neural excitability to influence the output of CeAm neurons. LA – lateral amygdala, BLA – basolateral amygdala.

### Experiment 2: Electrophysiological characterization of CeA CRF_1_:Cre -^td^Tomato neurons

#### Membrane properties and inhibitory transmission

^td^Tomato^+^ CRF_1_^+^ neurons were identified and differentiated from unlabeled CeA neurons using fluorescent optics and brief (<2 s) episcopic illumination in slices from adult male and female CRF_1_:Cre -^td^Tomato rats. Consistent with immunohistochemical studies (**Fig. 3**), the majority of CRF_1_^+^ neurons were observed in the medial subnucleus of the CeA (CeAm) and this region was targeted for recordings. Passive membrane properties were determined during online voltage-clamp recordings using a 10 mV pulse delivered after break-in and stabilization. The resting membrane potential was determined online after breaking into the cell using the zero current (I = 0) recording configuration. No differences were observed between male and female membrane properties including membrane capacitance, membrane resistance, decay time constant, or resting membrane potential (**Fig. 4A**). CRF_1_^+^ CeA neurons were then placed in current clamp configuration and a depolarizing step protocol was conducted to allow cell-typing based on previously-described firing properties (Chieng et al. 2006; Herman & Roberto, 2016). The majority of CRF_1_^+^ CeA neurons were of the low-threshold bursting type (**Fig. 4B**). Voltage-clamp recordings of pharmacologically-isolated GABA_A_ receptor-mediated spontaneous inhibitory postsynaptic currents (sIPSCs) revealed that CRF_1_^+^ neurons are under a significant amount of phasic inhibition (**Fig 4C**) with no significant sex differences in sIPSC frequency (**Fig 4D, left**) or sIPSC amplitude (**Fig 4D, right**).These data indicate that male and female CRF_1_^+^ CeA neurons have similar basal membrane properties and are under similar levels of basal inhibitory transmission.

**Figure 4.**
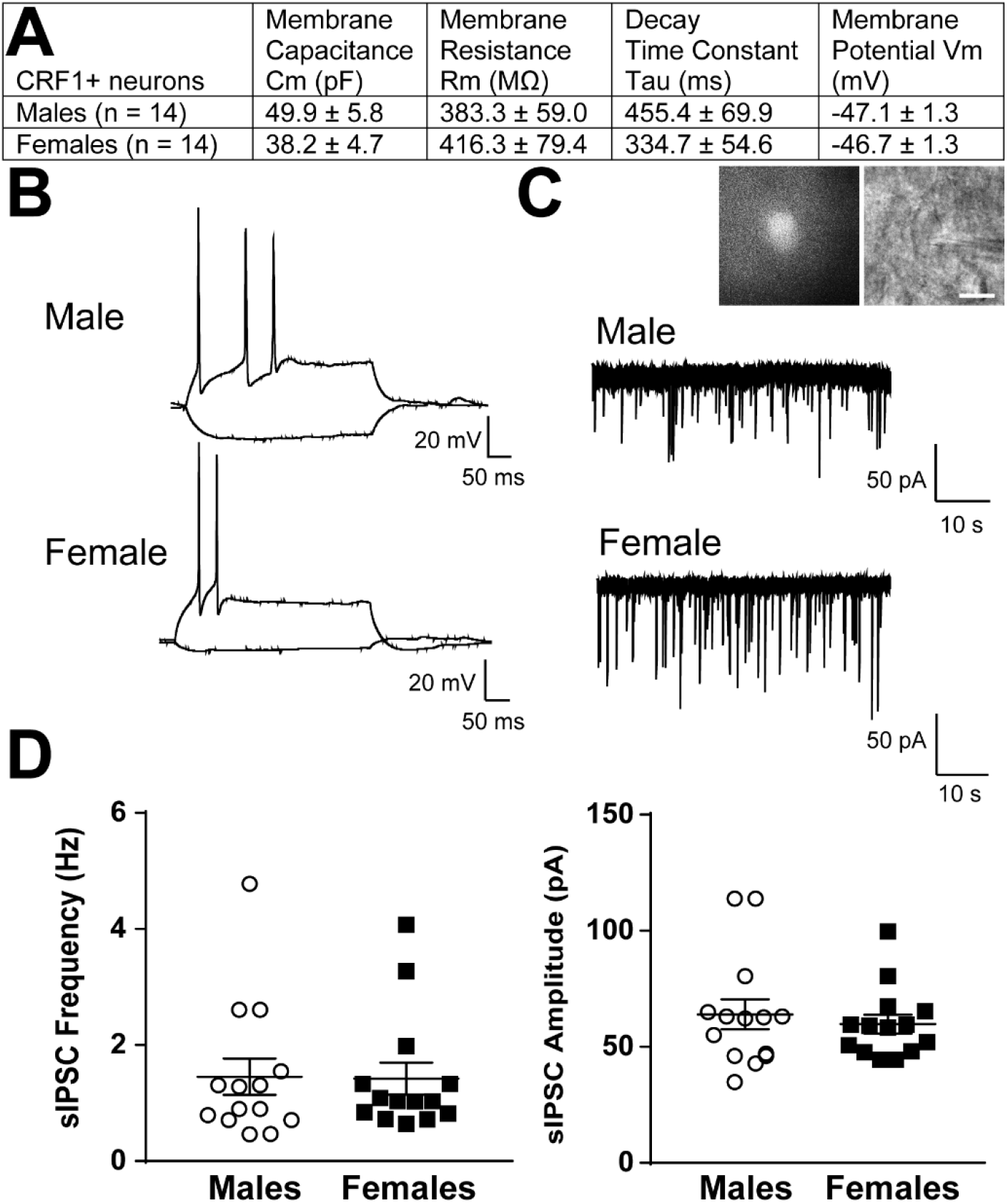
Basal membrane properties and inhibitory synaptic transmission in CeAm CRF_1_:Cre-^td^Tomato neurons. **(A)** Basal membrane properties (Membrane Capacitance, Cm; Membrane Resistance, Rm; Decay Constant, Tau; Membrane Potential, Vm) from male and female CRF_1_^+^ CeAm neurons. **(B)** Representative current-evoked spiking properties from male (top) and female (bottom) CRF_1_^+^ CeAm neurons. **(C)** Basal spontaneous inhibitory postsynaptic currents (sIPSCs) from male (top) and female (bottom) CRF_1_^+^ CeAm neurons (right). *Inset*: representative fluorescent (left) and infrared differential interference contract (IR-DIC, right) image of a CRF_1_^+^ CeAm neuron targeted for recording. Scale bar = 20 μm. **(D)** Average sIPSC frequency (left) and sIPSC amplitude (right) from male and female CRF_1_^+^ CeAm neurons.

#### CRF sensitivity

Spontaneous firing activity was recorded in CRF_1_^+^ CeA neurons from male and female CRF_1_:Cre-^td^Tomato rats using the cell-attached configuration. After a stable baseline period of regular firing was established, CRF (200 nM) was focally applied and the firing activity was recorded for a sustained application period of 7-12 min. CRF_1_^+^ CeA neurons from male rats had an average baseline firing rate of 2.1 ± 0.5 Hz and focal application of CRF significantly increased the firing activity to 3.4 ± 0.7 Hz (t = 3.5, p = 0.011; **Fig. 5A and 5C**). CRF_1_^+^ CeA neurons from female rats had an average baseline firing rate of 0.8 ± 0.2 Hz and focal application of CRF significantly increased the firing activity to 1.3 ± 0.3 Hz (t = 3.1, p = 0.016 by paired t-test; **Fig. 5B and 5D**). When firing activity was normalized to baseline values, CRF application significantly increased firing in CRF_1_^+^ CeA neurons from male rats to 192.3 ± 25.6 % of Control (t = 3.6, p = 0.009; **Fig. 5E**) and significantly increased firing in CRF_1_^+^ CeA neurons from female rats to 166.5 ± 13.9 % of Control (t = 4.8, p = 0.002; **Fig. 5E**) with no significant difference in the change in firing in response to CRF between male and female CRF_1_^+^ neurons.

**Figure 5.**
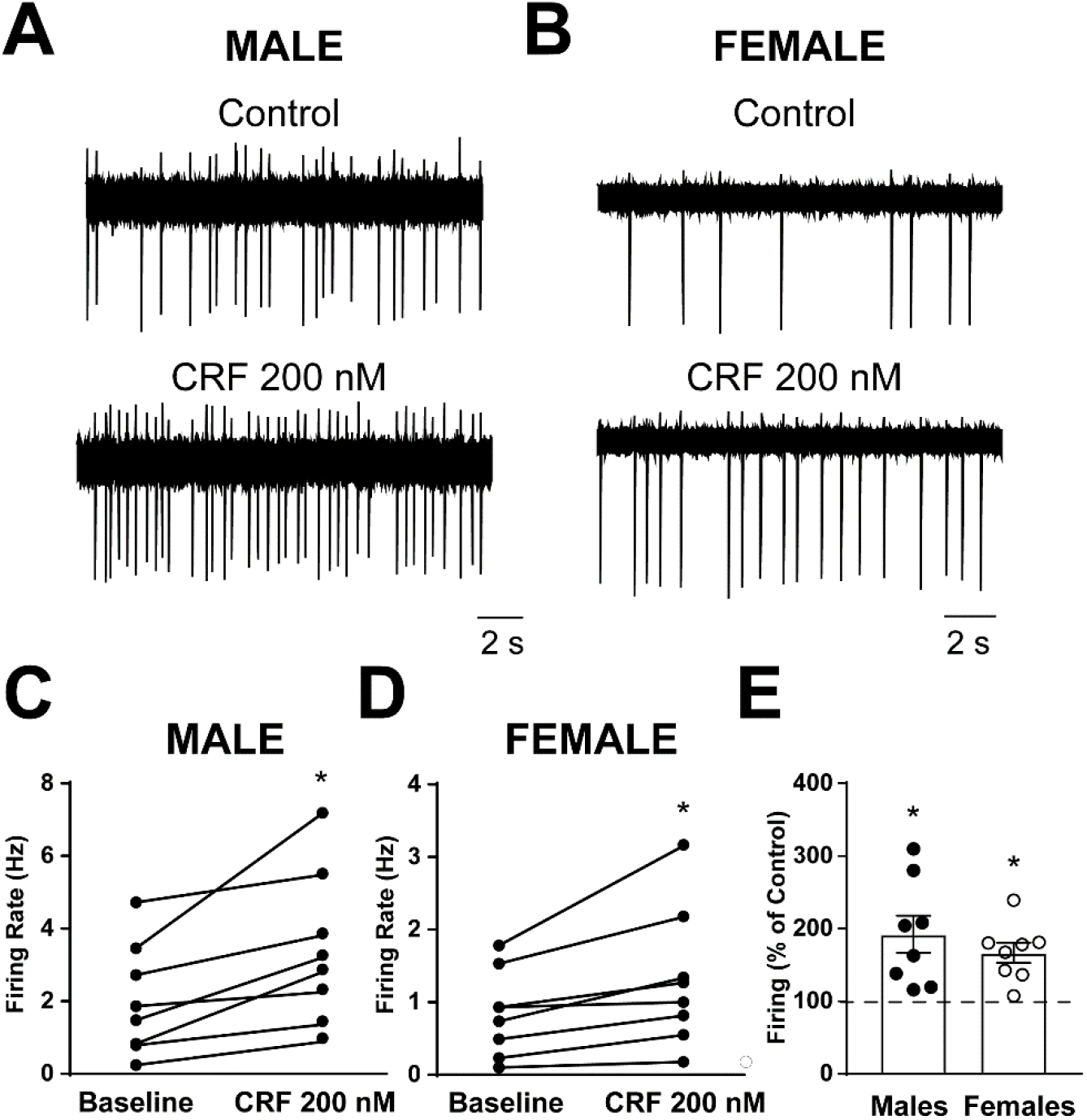
Spontaneous firing activity and CRF sensitivity of CeAm CRF_1_:Cre-^td^Tomato neurons. **(A)** Representative cell-attached recording of spontaneous firing activity in a CRF_1_^+^ CeAm neuron from a male CRF_1_:Cre-^td^Tomato rat before and during CRF (200 nM) application. **(B)** Representative cell-attached recording of spontaneous firing activity in a CRF_1_^+^ CeAm neuron from a female CRF_1_:Cre-^td^Tomato rat before and during CRF (200 nM) application. **(C)** Summary of changes in spontaneous firing activity with CRF application in CRF_1_^+^ CeAm neurons from male CRF_1_:Cre-^td^Tomato rats.*p<0.05 by paired t-test. **(D)** Summary of changes in spontaneous firing activity with CRF application in CRF_1_^+^ CeAm neurons from female CRF_1_:Cre-^td^Tomato rats.*p<0.05 by paired t-test. **(E)** Normalized change in firing activity in CRF_1_^+^ CeAm neurons from male and female CRF_1_:Cre-^td^Tomato rats.*p<0.05 by one-sample t-test.

### Experiment 3: Targeting of Cre-dependent Gq-DREADDs to the CeA increases c-Fos^+^ CRF_1_:Cre-^td^Tomato cells and CeA CRF_1_:Cre-^td^Tomato cell activity following CNO treatment

#### c-Fos immunohistochemistry

Four weeks after CRF_1_:Cre-^td^Tomato male and female rats were given intra-CeA microinjections of AAV8-hSyn-DIO-HA-hM3D(Gq)-IRES-mCitrine (Active Virus) or AAV5-hSyn-DIO-EGFP (Control Virus), rats were given an injection (i.p.) of CNO (4 mg/kg) and were sacrificed 90 min later. Brain sections containing the CeA were processed for c-Fos immunohistochemistry and the number of c-Fos^+ td^Tomato cells were quantified. An overwhelming majority of ^td^Tomato cells were located in the CeAm as shown above (**Fig. 3**). Therefore, quantification of c-Fos and ^td^Tomato cells was focused on this subregion. We found that rats in the active virus group had a higher percentage of c-Fos^+ td^Tomato cells than rats in the control virus group (*t* = 7.1, *p* < 0.001; **Fig. 6A, B**).

**Figure 6.**
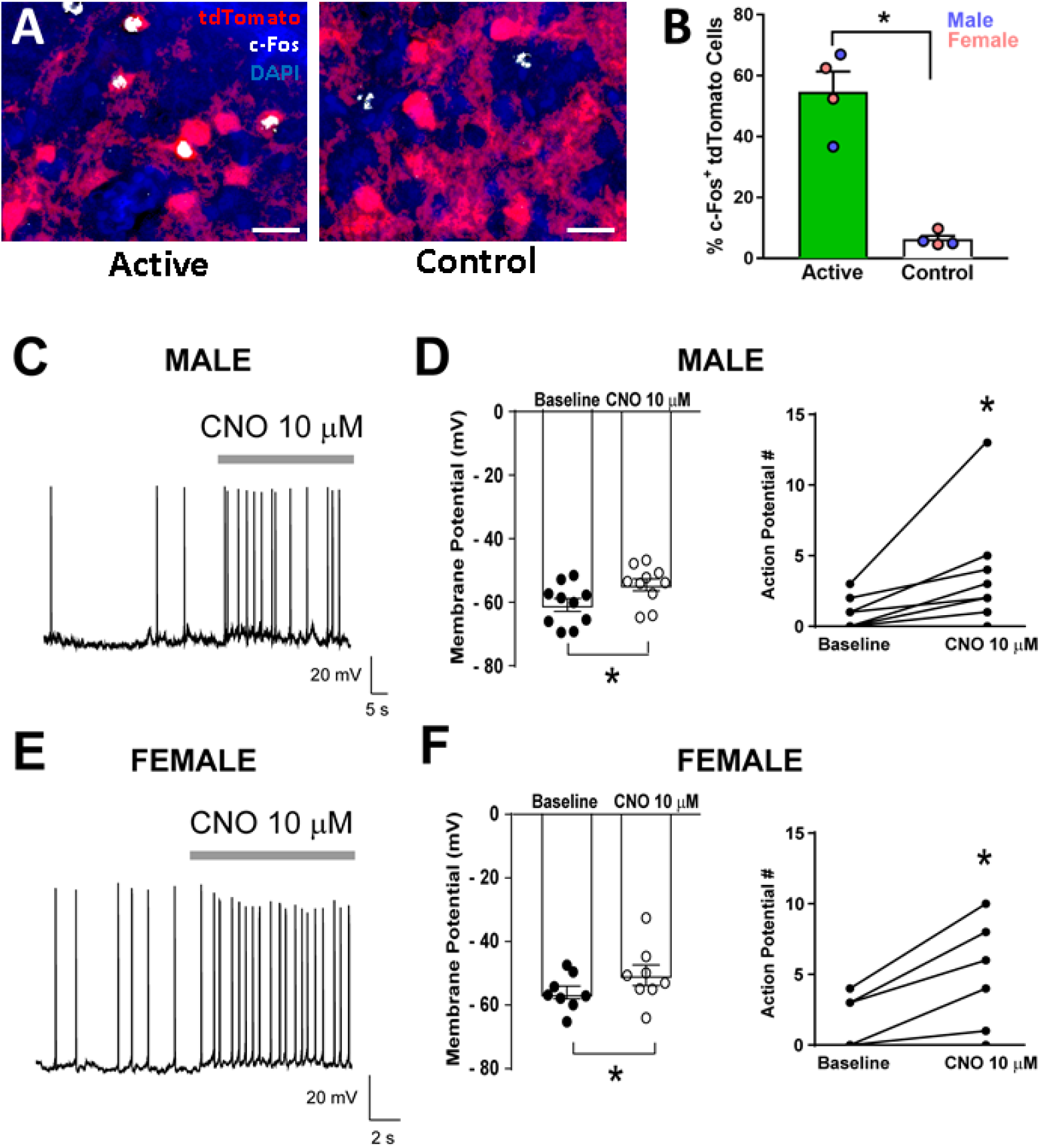
Validation of DREADD expression and function in CeA CRF_1_:Cre-^td^Tomato neurons. **(A)** Representative images of CRF_1_:Cre-^td^Tomato cells (red) and c-Fos immunostaining (white) in CeAm of rats that were given intra-CeA microinjections of AAV8-hSyn-DIO-HA-hM3D(Gq)-IRES-mCitrine (active virus) or AAV5-hSyn-DIO-EGFP (control virus). Scale bar: 50 μm. **(B)** CNO treatment 90 min before sacrifice increased the percentage of c-Fos^+ td^Tomato cells in CeAm of rats that were given active virus compared to rats that were given control virus microinjections. \**p* < 0.05. **(C)** Representative whole-cell current clamp recording of membrane potential and firing activity in a CRF_1_^+^ CeAm neuron from a male CRF_1_:Cre-^td^Tomato rat before and during CNO (10 μM) application. **(D)** Summary of the change in membrane potential (left) and action potentials (right) in male CRF_1_^+^ CeAm neurons after CNO application. *p<0.05 by paired t-test. **(E)** Representative whole-cell current clamp recording of membrane potential and firing activity in a CRF_1_^+^ CeAm neuron from a female CRF_1_:Cre-^td^Tomato rat before and during CNO (10 μM) application. **(D)** Summary of the change in membrane potential (left) and action potentials (right) in female CRF_1_^+^ CeAm neurons after CNO application. *p<0.05 by paired t-test.

#### Slice electrophysiology

Coronal sections containing the CeA were prepared from male and female CRF_1_:Cre-^td^Tomato rats >4 weeks after rats were given intra-CeA microinjections of AAV8-hSyn-DIO-HA-hM3D(Gq)-IRES-mCitrine. ^td^Tomato^+^ and mCitrine^+^ neurons were identified by brief episcopic illumination using fluorescent optics and positively-identified neurons were targeted for recording in whole-cell current clamp configuration to measure changes in resting membrane potential and spontaneous firing. CNO (10 μM) significantly increased membrane potential and number of action potentials in both male (t = 4.3, p = 0.002; t = 2.6, p = 0.030, respectively; **Fig. 6C** and **6D**) and in female CRF_1_^+^ mCitrine^+^ neurons in the CeA (t = 3.2, p = 0.016; t = 3.6, p = 0.006, respectively ; **Fig. 6E** and **6F)**, suggesting that hM3D(Gq) receptor expression was functional and could be stimulated by CNO application with no sex differences in expression or agonist sensitivity.

### Experiment 4: Chemogenetic stimulation of CeA CRF_1_:Cre-^td^Tomato cells increases mechanical nociception and anxiety-like behaviors

Rats were given intra-CeA microinjections of a viral vector for Cre-dependent expression of Gq-DREADD receptors (AAV8-hSyn-DIO-HA-hM3D(Gq)-IRES-mCitrine) or control fluorophore (AAV5-hSyn-DIO-EGFP) (**Fig. 7A**). Behavioral procedures began ≥4 weeks later (**Fig. 7B**).

**Figure 7.**
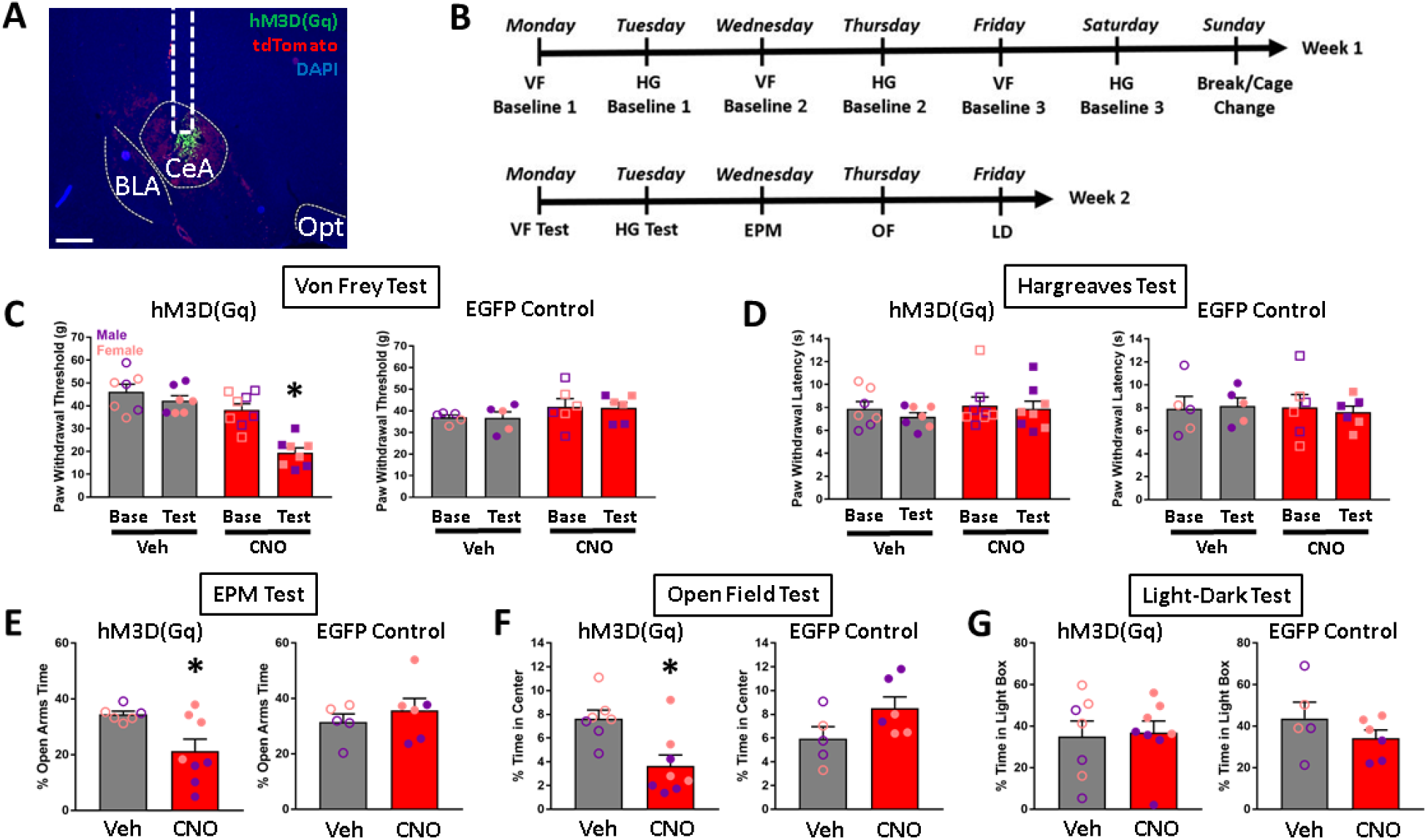
Effects of chemogenetic stimulation of CeA CRF_1_:Cre-^td^Tomato neurons on nociception and anxiety-like behaviors. **(A)** Representative image of AAV8-hSyn-DIO-HA-hM3D(Gq)-IRES-mCitrine expression (green) in the CeA. Scale bar: 500 μm. BLA: basolateral amygdala, Opt: optic tract. **(B)** Timeline of experimental procedures. **(C)** CNO treatment decreased paw withdrawal thresholds in the Von Frey test of mechanical nociception in rats that were given intra-CeA hM3D(Gq) virus microinjections. There were no effects of treatment on paw withdrawal thresholds in the EGFP control group. **(D)** CNO treatment had no effects on paw withdrawal latencies in either the hM3D(Gq) or EGFP groups in the Hargreaves test of thermal nociception. **(E)** CNO treatment decreased the percent time spent in open arms in the EPM test in the hM3D(Gq) group, but had no effect in the EGFP group. **(F)** CNO treatment decreased the percent time spent in the center of the arena in the OF test in the hM3D(Gq) group, but had no effect in the EGFP group. **(G)** CNO treatment had no effect on percent time spent in the light box in the LD test. \**p* < 0.05.

#### Nociception

In the Von Frey test of mechanical nociception, CNO treatment decreased paw withdrawal thresholds in rats that have hM3D(Gq) expression targeted to CeA CRF_1_^+^ cells [repeated measures ANOVA; test x treatment interaction (*F_1,13_* = 14.0, *p* = 0.002)]. There was a significant effect of test within the CNO group only (*F_1,7_* = 38.0, *p* < 0.001)], suggesting that chemogenetic stimulation of CeA CRF_1_^+^ cells increases mechanical sensitivity. CNO treatment had no effect on paw withdrawal thresholds in rats that received the control EGFP fluorophore (**Fig. 7C**). In the Hargreaves test of thermal nociception, CNO did not affect paw withdrawal latencies in either the hM3D(Gq) or control EGFP groups (**Fig. 7D**).

#### Anxiety-like behaviors

In the EPM test, in the hM3D(Gq) group, rats that were given CNO treatment had lower open arms time compared to rats that were given vehicle treatment (t-test; *t* = 2.6, *p* = 0.022), suggesting that chemogenetic stimulation of CeA CRF_1_^+^ cells increases anxiety-like behavior on the EPM. Control EGFP virus rats did not show differences in open arms times after CNO treatment (**Fig. 7E**). One rat in the hM3D(Gq) group that was given vehicle injection fell off the maze > 3 times and was therefore excluded from analysis. In the OF field test, hM3D(Gq) rats that were given CNO treatment spent less time in the center of the arena compared to rats that were given vehicle treatment (t-test; *t* = 3.3, *p* = 0.006), suggesting that chemogenetic stimulation of CeA CRF_1_^+^ cells increases anxiety-like behavior in the OF. EGFP control rats did not show differences in time spent in the center of the arena after CNO treatment (**Fig. 7F**). In the LD test, CNO treatment did not produce differences in time spent in the light box in either hM3D(Gq) or EGFP control groups (**Fig. 7G**). There were also no differences in latency to enter the light box (data not shown; 20.14 ± 5.63 s, 29.38 ± 10.42 s, 23.00 ± 3.94 s, and 16.17 ± 4.48 s, respectively, for rats in hM3D(Gq) – Veh, hM3D(Gq) – CNO, EGFP – Veh, and EGFP Virus – CNO groups).

### Experiment 5: iCre is expressed in CRF_1_-expressing neurons in BLA

The purpose of this experiment was to test the hypothesis that *Crhr1* and *iCre* mRNA are highly colocalized in cells in a brain region that contains a high density of glutamatergic neurons that express CRF_1_ receptors (i.e., the BLA). RNAscope ISH of brain sections containing BLA show dense *Crhr1* and *iCre* mRNA expression in BLA and surrounding areas (**Fig. 8A**). Quantification of *Crhr1* and *iCre* mRNA in the BLA showed that 98.5% of *Crhr1*-expressing cells co-express *iCre* (**Fig. 8B**, top), and that 99.4% of *iCre*-expressing cells co-express *Crhr1* (**Fig. 8B**, bottom).

**Figure 8.**
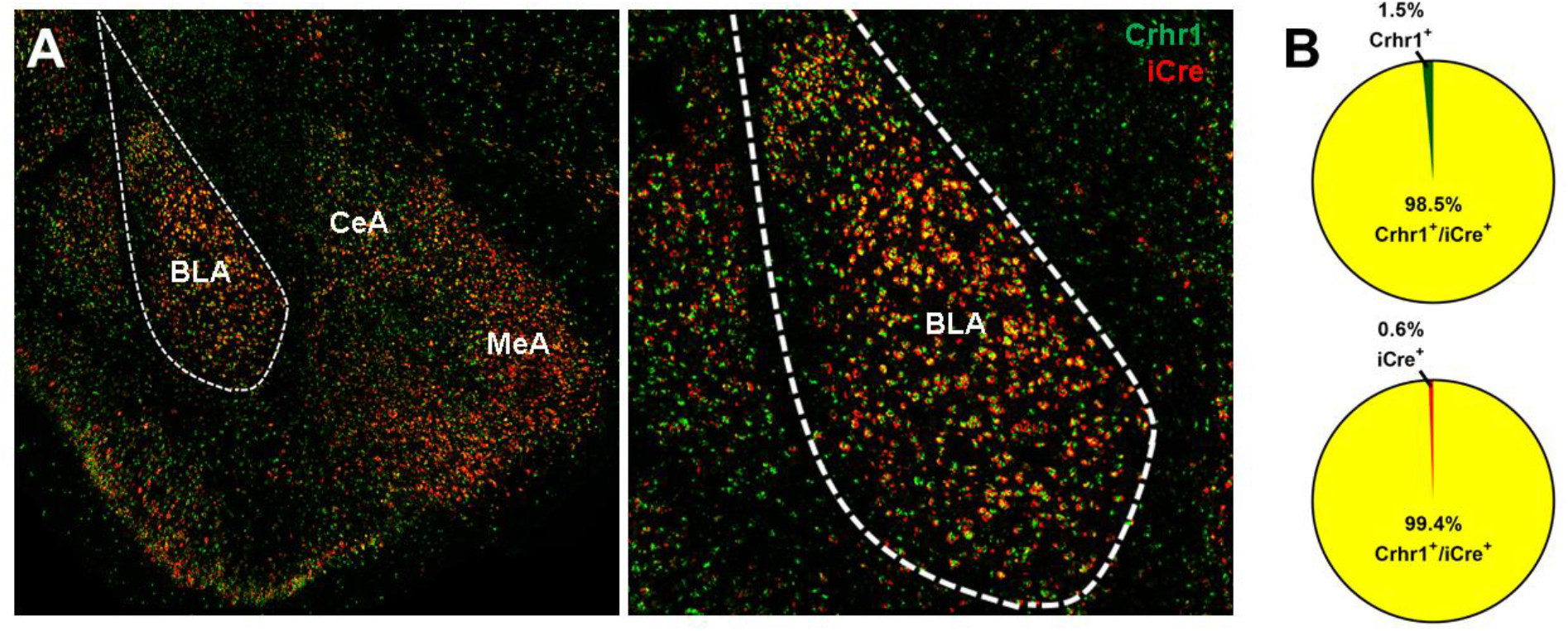
*Crhr1* and *iCre* mRNA expression in BLA of CRF_1_:Cre-^td^Tomato rats. **(A)** *Crhr1* (green) and *iCre* (red) mRNA are highly co-expressed in BLA and surrounding areas. **(B)** Within the BLA, 98.5% of *Crhr1^+^* cells co-express *iCre* (top), and 99.4% of *iCre^+^* cells co-express *Crhr1* (bottom).

## Discussion

We generated a new transgenic CRF_1_:Cre-^td^Tomato rat line to allow genetic manipulation and visualization of neurons that express CRF_1_ receptors in the rat brain. We report that, within the CeA of CRF_1_:Cre-^td^Tomato rats, CRF_1_:Cre-^td^Tomato cells are located in the medial subdivision (CeAm), consistent with previous reports of *Crhr1* expression in rats (Potter et al. 1994, van Pett et al., 2000; Day et al., 1999) and mice (van Pett et al, 2000; Justice et al., 2008), and that there is strong concordance (> 90%) between *Crhr1* and *iCre* mRNA expression in the CeAm of male and female CRF_1_:Cre-^td^Tomato rats. We also characterized the basal membrane properties, inhibitory synaptic transmission, and validated the CRF sensitivity of ^td^Tomato-expressing CeA CRF_1_^+^ cells in male and female rats. In addition, we showed that stimulatory DREADD receptors [hM3D(Gq)] can be targeted to CeA CRF_1_:Cre-^td^Tomato cells using a Cre-dependent expression strategy, that systemic CNO treatment induces c-Fos in hM3D(Gq)-transduced CRF_1_^+^ cells in CeA, and that CNO induces membrane depolarization and spontaneous firing activity in hM3D(Gq)-transduced CRF_1_^+^ cells in CeA. Finally, we showed that hM3D(Gq)-mediated stimulation of CeA CRF_1_^+^ cells increases anxiety-like behavior, as measured by EPM and open field tests, as well as mechanical nociception as measured by the Von Frey test. We also report that *Crhr1* and *iCre* mRNA expression are highly colocalized in the BLA. Collectively, this work provides cellular, electrophysiological, and behavioral data demonstrating the validity and reliability of a new transgenic rat model for the identification and selective manipulation of CRF_1_^+^ neurons in the CeA.

CRF_1_:Cre-^td^Tomato rats were generated by modifying a BAC clone derived from F344 rats, that was injected into oocytes derived from Wistar rats. Because the originating strain of the BAC DNA differs from the strain used for transgenesis, DNA sequence differences between F344 and Wistar rats in the 200+kb region surrounding the *Crhr1* genomic locus may impact expression of iCre-2A-^td^Tomato in the CRF_1_-Cre transgenic rat. Moreover, because transgenic BAC DNA insertion was not directed, position effect, copy number, and fragmentation of the BAC insertion may possibly alter expression of reporter genes. These sources of variability were minimized by the expertise of the UNC Transgenic Core Facility, where BAC DNA was prepared, cleaved at a single site to produce full-length single stranded BAC DNA, then injected into oocytes. Two transgenic founders were isolated that contained unique DNA sequences near the *Crhr1* genomic locus in the BAC, as well as DNA sequence from both terminal ends of the injected BAC, indicating insertion of full-length BAC DNA. Of these two rats, one displayed expression of ^td^Tomato in a pattern reflecting *Crhr1* expression patterns reported in rats (Potter et al., 1994; Pett et al., 2000), as well as transgenic expression of reporter genes reported in similarly designed BAC transgenic mice (Justice et al., 2008; Jiang et al, 2018; Hunt et al., 2018). The genomic locus and copy number of transgenic BAC insertion have not been further characterized. However, the high degree of alignment of Cre/^td^Tomato expression with known CRF_1_ expression patterns in the CeA, and broadly throughout the brain (manuscript in preparation), suggests that position, copy number, or fragmentation did not substantially alter expression of the transgene.

The intrinsic properties of CRF_1_^+^ (^td^Tomato^+^) CeA neurons in CRF_1_ :Cre-^td^Tomato rats are similar to those reported in wild-type rat CeA neurons (Herman & Roberto, 2016), suggesting that the transgene did not significantly impact the overall health or activity of CRF_1_^+^ CeA neurons. Previous studies have employed a CRFR1:GFP transgenic mouse model to examine CRF_1_^+^ CeA neurons in local CeA microcircuitry (Herman et al., 2013; 2016). Although prior work was conducted in mouse and some species differences would be expected, the electrophysiological properties of CRF_1_^+^ neurons in the CeA are relatively consistent between the mouse and rat transgenic models. Recent work specifically examining sex differences in CRF_1_^+^ neurons from CRFR1:GFP mice reported similar intrinsic membrane properties, cell-typing, and baseline inhibitory transmission as reported here and noted no sex differences in basal properties between male and female CRF_1_^+^ neurons in the CeA (Agoglia et al., 2020), consistent with our current findings. In contrast to previous work, however, we found no sex differences in CRF sensitivity of CRF_1_^+^ cells in CeA. Although CRF was previously found to increase firing in CRF_1_^+^ CeA neurons in both male and female mice, CRF_1_^+^ CeA neurons from male mice displayed a significantly greater increase in firing in response to CRF application (Agoglia et al., 2020). This discrepancy may be driven by differences in sampling size (although the current study was not sufficiently powered to detect sex differences), high variability in male baseline firing rates observed here, sex differences in baseline firing rates observed here, or it may reflect a species difference in sex-dependent sensitivity to CRF. Additional studies are required for a more comprehensive examination of sex- and/or species-specific differences in CRF-stimulated CRFR1^+^ neuronal activity in distinct brain regions. CRF_1_^+^ neurons in the CeA have also previously been implicated in the neuroplastic changes associated with acute and chronic ethanol exposure in mice (Herman et al., 2013; 2016), and future work will determine if this is also the case in rats.

To test the feasibility of using Cre-dependent stimulatory DREADDs to interrogate the role of CeA CRF_1_^+^ cells in behavior, we targeted Gq-coupled DREADD receptors [DIO-hM3D(Gq)] to CeA CRF_1_:Cre cells and first tested the effects of CeA CRF_1_^+^ cell stimulation on nociception. We showed that DREADD stimulation of CeA CRF_1_^+^ cells increases mechanical sensitivity as measured by the Von Frey test, but not thermal sensitivity measured by the Hargreaves test. Numerous studies have implicated CeA CRF-CRF_1_ signaling in nociception. For instance, pain induced by carrageenan injection into the knee joint produces hyperactivity of CeA neurons in rats, a phenomenon that is CRF_1_ dependent (Ji and Neugebauer, 2007). Conversely, latency for hind limb withdrawal reflex induced by knee joint pressure application is decreased (reflecting hyperalgesia) by CRF infusion into the CeA, a phenomenon that is blocked by co-infusion of a CRF_1_ antagonist (Ji et al., 2013). With regards to mechanical sensitivity, our finding that DREADD activation of CeA CRF_1_ neurons produces mechanical hypersensitivity extends previous work showing that ablation or inhibition of CeA CRF neurons blocks neuropathic pain-induced mechanical hypersensitivity, as measured by the Von Frey test (Andreoli et al., 2017). Various studies have also previously reported that CeA CRF_1_ antagonism attenuates mechanical hypersensitivity produced by chronic drug or alcohol exposure (e.g., Cohen et al., 2015; Edwards et al., 2012). Collectively, this prior work suggests that activation of CeA CRF-CRF_1_ signaling by chronic injuries or insults facilitates a hyperalgesic state, which was recapitulated in the current study via chemogenetic stimulation of CeA CRF_1_ cells. It should be noted that chemogenetic stimulation of CeA CRF^+^ cells has been shown to potentiate stress-induced analgesia (Andreoli et al., 2017), although this effect may be due to CRF release outside the CeA (eg., locus coeruleus) or due to changes in GABA release from CRF^+^ cells. Within the CeA, pharmacological studies suggest that pro- and anti-nociceptive effects of CRF signaling may be mediated by CRF_1_ and CRF2 receptors, respectively (Ji and Neugebauer 2007, 2008), but further studies are required to delineate the precise mechanisms by which CeA CRF signaling supports hyperalgesia and analgesia. The CRF_1_:Cre-^td^Tomato rat described here represents a useful tool for these studies.

Our lab has shown that predator odor stress-induced hyperalgesia, as measured by the Hargreaves test, is mediated by increased CRF-CRF_1_ signaling in the CeA (Itoga et al., 2016). Contrary to our hypothesis, we did not observe any effect of DREADD activation of CeA CRF_1_ cells on thermal sensitivity. It is not clear why CRF_1_:Cre rats exhibited mechanical hypersensitivity but not thermal hyperalgesia in this study. It is possible that specific CeA cell populations are involved in specific types of nociceptive processing, or it is possible that the engagement/recruitment of CeA CRF_1_^+^ cells in mediating specific types of nociception depends on the animal’s history or affective state. Using the CRF_1_:Cre-^td^Tomato rat line reported here, future work will elucidate the role of specific CRF_1_^+^ circuits in mechanical and thermal sensitivity (which may or may not be partially overlapping) under basal and challenged conditions such as neuropathic or inflammatory pain, stress exposure and/or withdrawal from chronic exposure to drugs or alcohol.

CeA CRF-CRF_1_ signaling generally promotes anxiogenic responses, particularly under challenged conditions such as stress or withdrawal from chronic exposure to drugs of abuse. For instance, in mice, intra-CeA infusion of a CRF_1_ antagonist attenuates anxiety-like behavior, as measured by open field and light-dark box tests, following immobilization stress but not under basal conditions (Henry et al., 2006). Similarly, blockade of CRF-CRF_1_ signaling in the CeA attenuates alcohol withdrawal-induced anxiety in rats, as measured by the EPM test (Rassnick et al., 1993). Exposure to stressors (Merlo-Pich et al., 1995) and alcohol withdrawal (Zorrilla et al., 2001) both increase extracellular CRF levels in the CeA. Messing and colleagues used CRF:Cre rats to show that CeA CRF cell activation (using DREADDs) increases anxiety-like behavior and that this effect is blocked by CeA CRF knockdown using RNA interference (Pomrenze et al., 2019). Collectively, these studies show that increased CRF-CRF_1_ signaling in CeA, whether produced by stress exposure, drug withdrawal, or the use of viral genetic tools, supports anxiogenesis. Here, we showed that chemogenetic stimulation of CeA CRF_1_^+^ cells increases anxiety-like behavior, as measured by EPM and open field tests. Interestingly, we did not observe an effect of chemogenetic stimulation of CeA CRF_1_^+^ cells on light-dark box measures, including time spent in the light vs. dark boxes, latency to enter light box, and number of crosses between the two compartments. Previous studies reported that the light-dark box test may be more suited for detecting anxiolytic rather than anxiogenic effects (Crawley and Davis, 1982), and that animals’ responses in this test is affected by the intensity of illumination in the animal’s regular housing room (File et al., 2004). Future studies will test if Cre-dependent expression of inhibitory chemogenetic or optogenetic strategies can be applied to inhibit CeA CRF_1_ cells, and if inhibition of CeA CRF_1_ cells reduces anxiety-like behavior at baseline or after stress exposure.

Although the CeA is generally not thought to exhibit strong sexual dimorphism, sex differences have been reported for CRF and CRF_1_ properties in the CeA. For example, female rats in proestrus have higher levels of CeA *Crf* mRNA than male rats and footshock stress induces greater CeA *Crf* expression in female proestrus than in male rats (Iwasaki-Sekino et al., 2009). However, using autoradiography, Cooke and colleagues reported no sex differences in CRF_1_ receptor binding in the CeA of rats (Weathington et al., 2014). Using a transgenic CRFR1:GFP mouse line, the CeA of male animals was shown to contain more CRF_1_^+^ cells than female animals (Agoglia et al., 2020). In the current study, although underpowered to detect sex effects, we found that male CRF_1_:Cre-^td^Tomato rats tend to have more CRF_1_^+^ cells within the CeA than female rats, as detected via RNAscope *in situ* hybridization. Previous work using CRFR1:GFP mice (Agoglia et al., 2020) and our current work using CRF_1_:Cre-^td^Tomato rats revealed no sex differences in basal membrane properties and inhibitory synaptic transmission in CeA CRF_1_^+^ neurons. Our previous work in CRFR1:GFP mice showed that, while CRF application increased firing in CRF_1_^+^ CeA neurons from both female and male mice, larger increases were observed in CRF_1_^+^ neurons from male mice, an effect that we did not observe in rats potentially due to differences in variability or sex differences in firing rate.

The overwhelming majority of published studies examining the role of CeA CRF_1_ signaling in nocifensive and anxiety-like behaviors has employed only male subjects. Here, we used both sexes in tests of nociception and anxiety-like behavior and we did not detect any sex differences. However, we observed that the anxiogenic effect of CeA CRF_1_^+^ cell activation in the EPM test was driven by stronger effects in male subjects. One interpretation is that CeA CRF_1_^+^ cell activation affects specific components of anxiety-related behavior, that these components are aligned with specific “anxiety-like behavior” phenotypes in male versus female rats that are more or less detectable by these various tests. In support of this idea, a meta-analysis of studies of anxiety-like behavior in rats and mice revealed a discordance in results between the EPM and open field tests (Mohammad et al., 2016). However, due to the small number of studies testing female animals, the role of sex in this discordance is still unknown. Within the EPM literature, studies have shown that the EPM test is more reliable for detecting anxiogenic effects in male rather than female rodents (e.g., Scholl et al., 2019). Collectively, these findings demonstrate the importance of behavioral assay selection and of using both male and female subjects.

In summary, we present a novel CRF_1_:Cre-^td^Tomato rat line that allows for the visualization and manipulation of CRF_1_-expressing cells. The CRF_1_:Cre rat can be used to study the role of distinct CRF_1_^+^ cell populations and circuits in behavior. In addition, the expression of the fluorescent reporter ^td^Tomato in CRF_1_-expressing cells will allow for high resolution, detailed analysis of the expression pattern of CRF_1_ that has not been possible due to the lack of CRF_1_-specific antibodies. Subsequent comprehensive validation studies will report the accuracy of CRF_1_ and Cre-^td^Tomato co-expression throughout the rat brain, and this transgenic line will be useful for anatomical, electrophysiological and functional analysis of CRF-CRF_1_ neural circuits in rats.

## Acknowledgements

This work was supported by National Institutes of Health Grants AA023305 (NWG), AA026022 (to NWG and MAH), AA023002 (MAH), AA027145 (MMW), and AA007577 (institutional NRSA training grant that supported MMW). This work was also supported in part by a Merit Review Award #I01 BX003451 (to NWG) from the United States Department of Veterans Affairs, Biomedical Laboratory Research and Development Service.

